# Improving Multi-Trait Genomic Prediction Efficiency Through The Incorporation Of Synthetic Traits Selected Based on Co-heritability

**DOI:** 10.1101/2025.06.23.661178

**Authors:** Ashmita Upadhyay, Ruhana Azam, Meilu Yuan, Stefan Ivanovic, John N. Ferguson, Rachel E. Paul, Oluwasanmi Koyejo, Mohammed El-Kebir, Alexander E. Lipka, Andrew D. B. Leakey, Samuel B. Fernandes

## Abstract

Genomic prediction (GP) is an essential tool in the field of plant breeding to accelerate the cultivar development pipeline by predicting the performance of unphenotyped lines. The precision of prediction is constrained by the heritability of the target trait when applying a single-trait genomic prediction model. To overcome this limitation, a multi-trait genomic prediction model leveraging high-heritability secondary traits co-heritable with the target trait can boost predictive ability for the target trait. However, this is practically challenging because it requires additional phenotyping effort and prior knowledge of trait co-heritability. This study aimed to assess the efficiency of multi-trait genomic prediction models powered by secondary traits derived from high-throughput phenotyping data when predicting important leaf functional target traits, i.e., nitrogen (N) content and specific leaf area (SLA) in diverse sorghum accessions. Since these traits can be predicted from hyperspectral reflectance data, there is significant potential for other wavelengths within the existing dataset to meet the criteria needed to improve prediction accuracy using multi-trait approaches. Therefore, experiments were performed on traditional direct measures of leaf N content and SLA, plus partial least squares regression predictions of them (Leaf N-PLSR, SLA-PLSR), i.e., four target traits in total. Three secondary, “synthetic traits” (S1, S2, S3), each a ratio of two wavelengths within the hyperspectral data, were identified based on high co-heritability with a given target trait. Single-trait GBLUP (Genomic Best Linear Unbiased Predictor) was fitted as a baseline model, followed by three multi-trait GBLUP models using synthetic traits and target traits together. Model performance was assessed using k-fold (k=5) cross-validation (CV), which consisted of single-trait, CV1, and CV2 schemes. The synthetic traits’ high genetic correlation and heritability met the requirements for their use as secondary traits. There was a significant increase in accuracy when synthetic traits were used in the multi-trait genomic prediction model compared to a single trait alone for all four target traits. It improved prediction accuracy while using secondary traits derived from hyperspectral high-throughput phenotyping data in the multi-trait genomic prediction model, suggesting that this approach could be broadly applied in a post-hoc fashion to many datasets without any additional phenotyping effort. Our analysis highlights a practical approach to improve multi-trait genomic prediction model performance using synthetic traits with no intrinsic biological meaning selected through co-heritability estimation.

## 1 Introduction

Genomic prediction (GP) is an essential tool in plant breeding that reduces the time required to release a new cultivar. It thus contributes to improving resilient food, fuel, bioproducts, feed, and fiber production. The strength of GP stems from its ability to utilize the phenotypic and genotypic information from a training population to predict the performance of individuals that were not evaluated in field trials [27], thereby increasing selection intensity in a breeding cycle [58]. Examples of successful applications of GP include prediction of disease resistance in wheat [3], nitrogen concentration in corn [6], yield prediction in wheat [8, 14], soybean [15], and rice [24], and biomass in sorghum [19]. However, the accuracy of prediction is bounded by the square root of the heritability 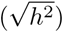 [11, 25, 26, 29], limiting the usefulness of GP for traits of low *h*^2^.

One approach that has been used to improve prediction accuracy in traits of low *h*^2^ is multi-trait GP (MT) [19–21, 29, 31]. Including a secondary trait in the model improves predictive ability when the secondary trait has a high *h*^2^ and is highly genetically correlated with the target trait [11, 25, 26, 29]. However, MT can be practically challenging due to the need for additional data collection and prior knowledge of a secondary trait with high heritability and correlation with the target trait. In the past, breeding programs were limited to phenotyping a few agronomic or morphological measures. However, with recent advances in remote sensing, robotics, and artificial intelligence-enabled analysis, it is now possible for a plethora of traits to be easily collected with high-throughput phenotyping (HTP) methods. Furthermore, there is commonly a lot of unused information in hyperspectral data. This study tests whether this untapped information in HTP data can be leveraged to ease the adoption of MT.

One commonly used modality of HTP data in plant science is hyperspectral reflectance spectra, which describe the intensity of reflected light at hundreds or thousands of wavelengths in the visible, near-infrared, and short-wave infrared portions of the electromagnetic spectrum [34]. In many cases, a subset of wavebands are used to estimate some of the many vegetation indices that have been previously defined as useful proxies of vegetation form and function e.g., normalized difference vegetation index (NDVI) [2, 15], soil adjusted vegetation indices (SAVI) [43], or NDRE. Alternatively, partial least squares regression models can be built to predict traits of interest from spectral data [55, 59]. Whichever approach is taken, there is commonly a significant amount of potentially useful information in the spectra that is collected but not utilized.

While GP and HTP data are independently powerful approaches, the optimum scenario would be an efficient integration of both tools. Attempts at this task have included obtaining a relationship matrix based on the HTP data [35] and utilizing previously validated indices as secondary traits in MT models [50]. While improvements were observed in the former, in the latter, there is a bottleneck where it is necessary to validate that the index selected possesses the characteristics of a favorable secondary trait. As the requirement for secondary traits to be beneficial in MT revolves around correlation with the target trait as well as heritability, one potential option to summarise these measures is to calculate the co-heritability (co*h*^2^) [17, 54]. Therefore, we hypothesize that the *coh*^2^ can be utilized to identify traits within HTP data that were not selected a priori or that inherently have intrinsic biological value but are powerful and flexible secondary traits for use in GP.

Hypothesis testing in this study uses leaf nitrogen content per unit area (Narea) and specific leaf area (SLA) of field-grown sorghum as a case study. Monitoring leaf nitrogen in the field is important for understanding many processes, including leaf metabolism, growth, ecosystem function, herbivory, litter decomposition, plant-microbe interactions, and fertilizer management (Melillo et al. 1982; Muchow and Sinclair 1994; Poorter and Evans 1998; Hamilton et al. 2001; Diaz et al. 2004; Uribelarrea et al. 2009; Sayer et al. 2017). Specific leaf area (SLA) – leaf lamina area per unit dry mass – is a key leaf trait tightly linked to variation in leaf composition, physiology, plant growth strategy, and resource investment (Shipley et al., 2006; Flores et al., 2014). Leaves with a high SLA typically have less nitrogen per unit leaf area, lower rates of photosynthesis, lower rates of respiration, are less well defended against herbivory, and decompose more rapidly than leaves with a low SLA (Wright et al., 2004).

## 2 Materials and method

In 2017, 869 photoperiod-sensitive sorghum lines were grown in two locations on the University of Illinois at Urbana-Champaign research farm, as described previously [16, 18, 44]. The experimental layout of each location was an augmented block design, with 16 incomplete blocks, and six check lines were included in each block. Each replicate field comprised 960 four-row plots of 3 m length, 1.5 m alley, and 0.76 m row spacing. Phenotyping was performed on the youngest fully expanded leaves of two randomly selected plants from the middle two rows of each plot as part of the sampling protocol described by [18]. The leaves were excised just above the ligule at dawn, and the cut end was immediately placed in water. In the laboratory, hyperspectral reflectance was measured at six locations on the adaxial surface in the central section of each leaf using a full-range spectroradiometer (ASD FieldSpec4® Standard-Res, Analytical Spectral Devices, Boulder, CO) equipped with an illuminated leaf clip. To conform with experiments on other species with smaller leaves, the leaf clip was fitted with a custom-made mask to limit illumination to an area of 150 mm2. A reflective white reference disc (Spectralon, Labsphere Inc., North Dutton, NH) was used to standardize the relative reflectance of the samples. The instrument was turned on 30-60 minutes prior to use to allow the three arrays to warm up. Each spectrum collected was the internal average of 10 scans of the instrument, each taking 100ms. Before averaging the measurements for a single leaf, a splice correction was applied to each spectrum to correct for the difference between the three sensors, and quality control was applied using the FieldSpectra package in R [49]. Subsequently, three 1.6 cm2 leaf discs were collected from the same portion of the leaf where hyperspectral reflectance was assessed. Leaf discs of known area were immediately transferred to an oven set at 60 °C for two weeks. SLA was calculated as the ratio of leaf area to dry mass. Leaf N content was determined using an elemental analyzer as described by [23] after the leaf discs were pooled and powdered using a mechanical grinder (2000 Grinder, SPEX CertiPrep, Metuchen, NJ, USA). Partial least squares regression (PLSR) models were built to predict Narea and SLA [44]. These PLSR-based predictions of Narea (PLSR-Narea) and SLA (PLSR-SLA) were used alongside traditional measures of Narea and SLA as the four target traits in this study.

### 2.0.1 Heritability estimation

The overarching hypothesis was that secondary traits derived from hyperspectral data could boost the predictive ability of target traits in multi-trait genomic prediction. Consequently, we sought to identify ratios of wavelengths within spectra with i) the highest heritability and ii) a high genetic correlation with the target trait of interest, i.e., Narea, SLR, PLSR-Narea, and PLSR-SLA. We focused on these two conditions because the resulting wavelength ratios exhibiting these properties were theoretically best suited for multi-trait models [41]. Accordingly, the first step towards identifying these best-suited ratios was to estimate the heritability of the traits collected from these two locations. For that, we used the following linear mixed model:

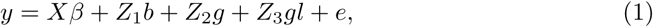

where *y* is the vector of phenotypes; *X* is the incidence matrix associated with the vector of fixed effects *β*, which includes set and location; *Z*_1_ is the incidence matrix associated with the vector of random effect of block *b* within set *s* and location *l*, with 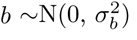; *Z*_2_ is the incidence matrix associated with the vector of random effect of genotypes *g*, with 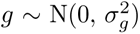; *Z*_3_ is the incidence matrix associated with the vector of random effect of genotypes by locations interaction, with 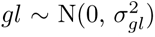; and *e* is the vector of residuals, with 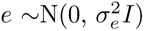. The terms 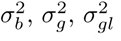, and 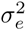 are the variances of block within the set and location, genotypes, genotypes by locations, and residuals. Finally, *I* denotes the identity matrix. Model 1 was fitted with the package LME4 [7] on the software R 4.3 (R Core Team). Model 1 was used to estimate variance components for the four target traits evaluated (Narea, SLA, PLSR-Narea, PLSR-SLA). Because individual wavelengths are not as informative, we investigated ratios of pairs of wavelengths (e.g., 400 nm / 2100 nm) [32, 37, 51, 56]. Thus, utilizing the 2150 individual wavelengths, we obtained 4,620,350 pairwise ratios. Once the variance components were obtained for the target traits and the wavelength ratios, the sample heritability (*ĥ*^2^) was estimated using the equation below:

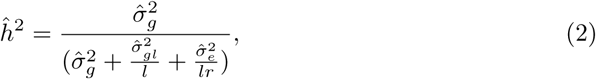

in which *l* is the number of locations, *r* is the harmonic mean of the number of replications, and all other terms are estimates of the parameters defined in model 1. Because this stage involved fitting millions of models, we used a simple heritability estimator, but more robust ones are available [45].

### 2.0.2 Co-heritability estimation and identification of synthetic traits

We next sought to determine which particular wavelength ratios were best suited to include as additional response variables in multi-trait genomic prediction. Such ratios were called synthetic traits, and these exhibited the strongest co-heritabilities with the target trait. The co-heritability between two traits is defined as their heritable genetic correlation and was calculated as follows [17]:

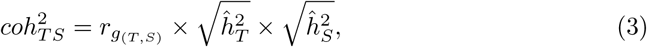

where 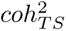 is the co-heritability between the target and the synthetic (i.e., secondary) trait, 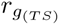 is the genetic correlation between the target and the synthetic trait, and 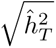 and 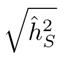 are the square root of sample heritability for the target trait and synthetic trait, respectively. To estimate the genetic correlation, we utilized the property of the variance of the sum of two random variables. First, we fitted a mixed linear model (1) to estimate the genetic variance for the target trait (i.e., Narea, SLA, PLSR-Narea, or PLSR-SLA), which was referred to as *σ_g_* in model 1, but to be more clearly associated with the target we will call it *σ_T_* and the genetic variance for the wavelength ratio will be called *σ_S_* to be associated with the synthetic trait. Second, we estimated the variance component for the target trait plus the wavelength ratio (e.g., *N* + (*Wave*_1000_*/Wave*_200_)). In this case, we estimated the genetic covariance of the two variables:

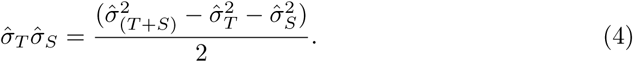

Using these estimates of genetic variances and covariance, we next estimated the genetic correlation between *T* and *S* as follows:

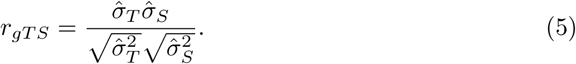

Because the genetic correlation can be positive or negative, to simplify co-heritability selection, we took the absolute value of *r_gT S_* (i.e., *r_gT S_*) for further use in the co-heritability estimation. Combined with the sample heritabilities of *T* and *S* calculated using 2 (denoted respectively as 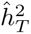 and 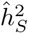), we then calculated the co-heritabilty with equation 3.

For each of the four target traits, 4,310,175 co-heritability values were estimated, one for each wavelength ratio. Heatmaps for each trait were generated to visualize the co-heritability across all traits, which can be found in Supplementary file 1 (see Appendix). To select a manageable number of synthetic traits for multi-trait genomic prediction, we selected the 1% wavelength ratios with the greatest co-heritability. Next, we calculated the Pearson correlation and Euclidean distance between these selected wavelength ratios. These Euclidean distances were subsequently used in a hierarchical cluster analysis to determine the extent to which these selected wavelength ratios clustered together. Three synthetic traits were selected from each group in a hierarchical cluster analysis. Ultimately, this analysis identified three representative wavelength ratios for use as synthetic traits in multi-trait genomic prediction. This procedure was repeated once for each target trait (i.e., Narea, SLA, PLSR-Narea, and PLSR-SLA). For each target trait, a synthetic trait with co-heritability of zero was also selected as a control against which to assess any possible improvement in performance when utilizing highly co-heritable traits in the multi-trait framework.

### 2.1 Genotypic data

The genotypic data set used in this study was previously published in [18]. Briefly, a set of 100,435 SNPs from [16] was imputed (Beagle 4.1) using the whole-genome resequencing data set of 5,512,653 SNPs published in [52] as a reference. The resulting data set consisted of 5.5 million SNPs. This data set was filtered to remove SNPs with low imputation quality (*R*^2^ *<* 0.5), markers with minor allele count less than 20 (vcftools), and SNPs with linkage disequilibrium higher than 0.9 in a window of 50 SNPs (Plink). This resulted in a total of 454,393 markers scored in 836 sorghum lines, which were considered in the ensuing analyses.

#### 2.1.1 Genomic Prediction

Genomic prediction was implemented in two stages. First, for each location, we obtained the best linear unbiased estimators (BLUEs) for all target and synthetic traits using linear mixed models implemented in ASReml-R [10]. The statistical model for BLUEs estimation was as follows:

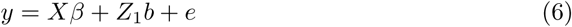

*y* is the phenotypic observation of selected synthetic and target trait; *X* is the incidence matrix associated with the vector of fixed effect *β*, which includes genotype *g* and set *s*; *Z*_1_ is the incidence matrix associated with the vector of random effect of block *b* with 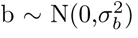; and *e_ijk_* is the random effect of residuals, with 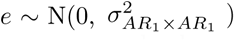, where *AR*_1_ × *AR*_1_ is a first-order auto-regressive structure applied to rows and columns for spatial correction.

Next, we fitted the following genomic prediction model as a second-stage model using BLUEs from the first stage for a single trait:

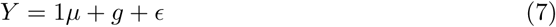

where Y is an *n* × *t* matrix of BLUEs of genotype effects from Model 6 estimated for each of t traits across n individuals; *µ* is a t-dimensional vector of grand mean parameter values for each of t traits, *g* is an *n* × *t* matrix of genetic effects for each of t traits following a matrix-variate normal distribution denoted 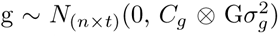 with *C_g_* denoting the genetic variance-covariance between traits, *G* denoting the additive genetic relatedness matrix [53], and 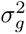 denoting the genetic variance parameter; and *e* is an *n* × *t* matrix of error terms 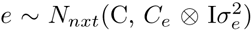 with *C_e_* denoting the non-genetic variance-covariance between traits, *I* denoting the identity matrix, and 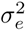 denoting the error variance parameter. In this study, n = 822 sorghum lines with genotypic information, t = 1 for the single-trait model that included only one of the four target traits, and t = 2 for the models that included one of the four target traits and one of the corresponding synthetic traits.

### 2.2 Cross-validation scheme

The usefulness of the synthetic traits to boost predictive ability in genomic prediction was assessed through *k*-fold cross-validation (CV), with *k* = 5 [33] with 20 repetitions, where each repetition has 5-fold cross-validation. Following the procedure described in [19], where three distinct scenarios were considered:

#### 2.2.1 Single-trait(ST)

Phenotypic values of a single target trait on 80 % of the genotypes were used for training a single-trait GBLUP model to predict genomic estimated breeding values (GEBVs) of the target trait for the remaining 20% of individuals for validation. Here, a single-trait GBLUP model, described in (8), was fitted in the training set with the target trait as the response variable. Predictive ability was evaluated using two metrics: the Pearson correlation coefficient between the observed target trait value and the corresponding GEBV from the single-trait GBLUP model fitted in the training set.

#### 2.2.2 CV1

Phenotypic data on the target trait and one of the corresponding synthetic traits from 80% of the genotypes were used to predict the GEBVs of the target trait for the remaining 20% of the genotypes. In this scenario, the multi-trait GBLUP model, described in (8), was fitted in the training set with the target trait and one of the synthetic traits as the two response variables. Predictive ability was evaluated using the same two metrics described in the ST scenario.

#### 2.2.3 CV2

Phenotypic data on the target trait from 80% of the individuals and phenotypic data on one of the corresponding synthetic traits from all 100% of genotypes were used to predict the GEBVs of the target trait for the remaining 20% of genotypes. Here, the multi-trait GBLUP model, described in model 7, was fitted to all genotypes for the target trait and the synthetic trait in the training set, as well as to all genotypes for the synthetic trait in the validation set. The same metric described in the ST scenario was used to evaluate predictive ability.

The same folds were used to assess performance across the three scenarios to enable a thorough comparison of prediction accuracies. To minimize the possibility of overfitting and obtain an over-optimistic assessment of model performance, as described in [47], we used one location for training and another location for validation. For instance, if the prediction model was trained on phenotypic information from genotypes evaluated in location EF, the validation set would use phenotypic information from genotypes evaluated in location MW and vice versa.

### 2.3 Control measure

A control measure was taken to validate the idea of selecting the synthetic trait that was highly heritable and genetically correlated with the target trait. The opposite was also done, where one synthetic trait with a co-heritability value of zero was selected for each target trait and was further used to fit the multi-trait genomic prediction model, where the model performance was assessed using CV1 and CV2 schemes.

### 2.4 20:80 approach

We conducted an additional analysis to explore the extent to which using the same individuals to identify synthetic traits increased the predictive ability of our multi-trait GBLUP models. Specifically, a testing framework was developed to identify synthetic traits on a subset of 20% of the individuals. In contrast, the remaining 80% were utilized within the cross-validation context as described in Figure 2. Thus, 20% of the individuals were randomly selected to generate three synthetic traits as described in equation 3. Next, the dataset containing the remaining 80% individuals was used to fit both single-trait and multi-trait genomic prediction models. This testing process was repeated five times (subsets one to five) to minimize bias. Model performance was assessed using the cross-validation schemes CV1, CV2, and the one described in ST. The predictive ability of models fitted using this 20:80 sampling approach was then compared to that of models in which the synthetic traits were obtained using the entire data.

**Figure 1.**
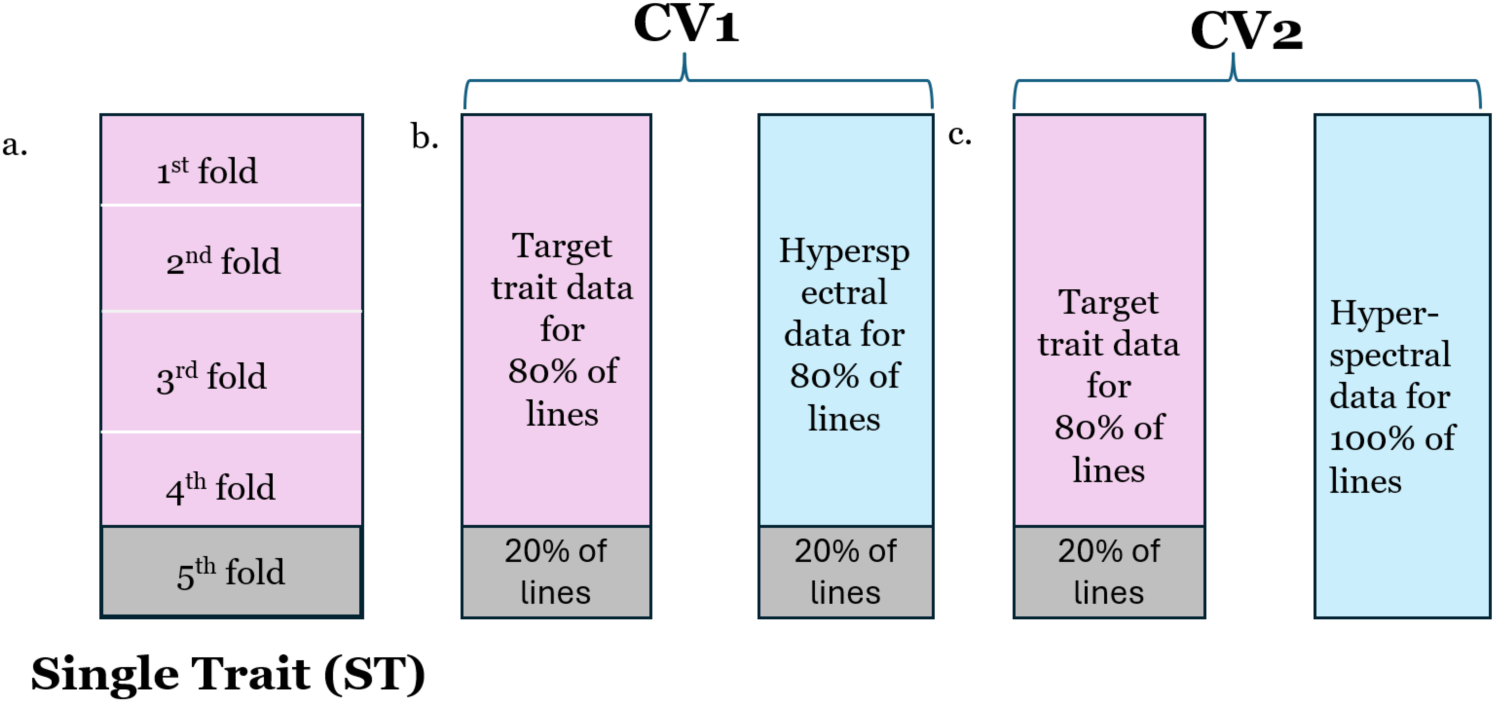
Diagrammatic representation of cross-validation scheme performed in single-trait and multi-trait genomic prediction model where only target trait’s phenotypic data is used in ST scheme, however, both phenotypic data, i.e., target trait and hyperspectral data, are used for CV1 and CV2

**Figure 2.**
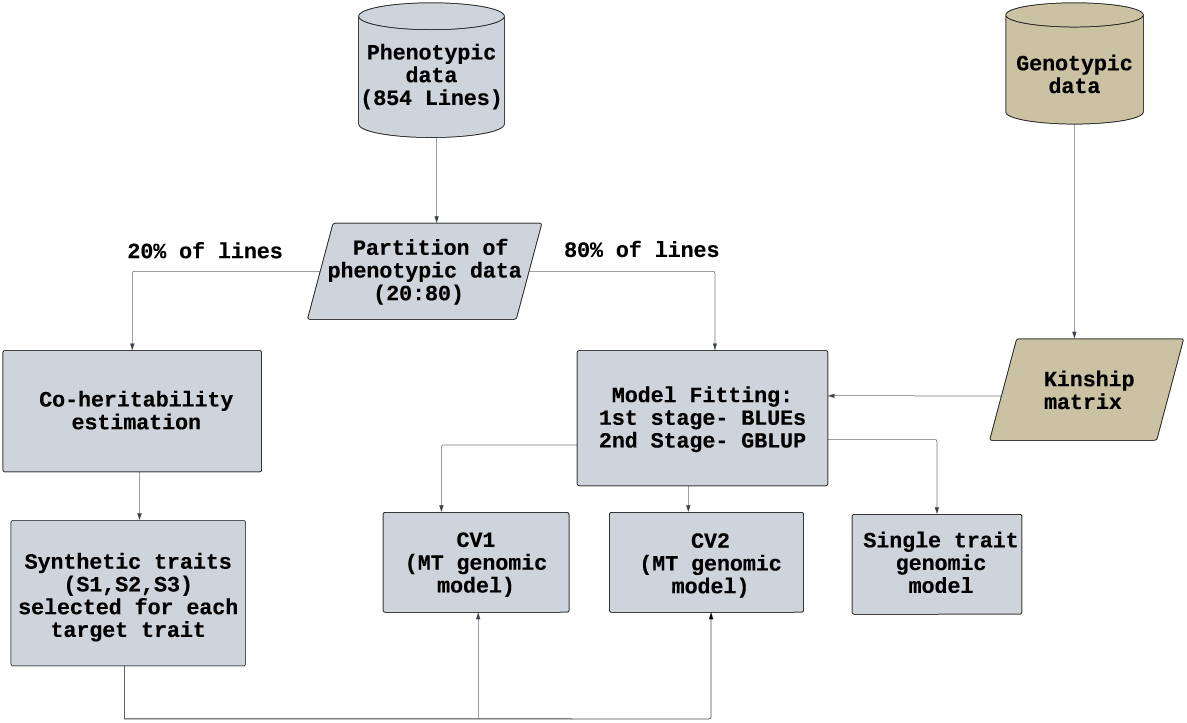
Flow chart with the methods used to separate 20% of the individuals from the whole data, estimate the co-heritability, select three synthetic traits, and use the selected synthetic trait further in a multi-trait (MT) genomic prediction model fitted for 80% of the data for four target traits

## 3 Results

Four target traits were used as case studies, and for each one, we identified three synthetic traits (S1, S2, S3) to be included in the multi-trait genomic prediction models. Thus, we identified 12 different synthetic traits. The target traits utilized here were only moderately heritable with *h*^2^ of 0.34, 0.32, 0.4, and 0.26 for Narea, SLA, PLSR-Narea, and PLSR-SLA, respectively (Figure 3). Conversely, the synthetic traits had higher heritability, where *h*^2^ ranged from 0.61 to 0.68. Also, the genetic correlation (*cor_g_*) of the synthetic traits with the respective target traits was high, ranging from 0.72 to 0.99. Finally, the co-*h*^2^ for the 12 synthetic traits ranged from 0.46 to 0.57.

**Figure 3.**
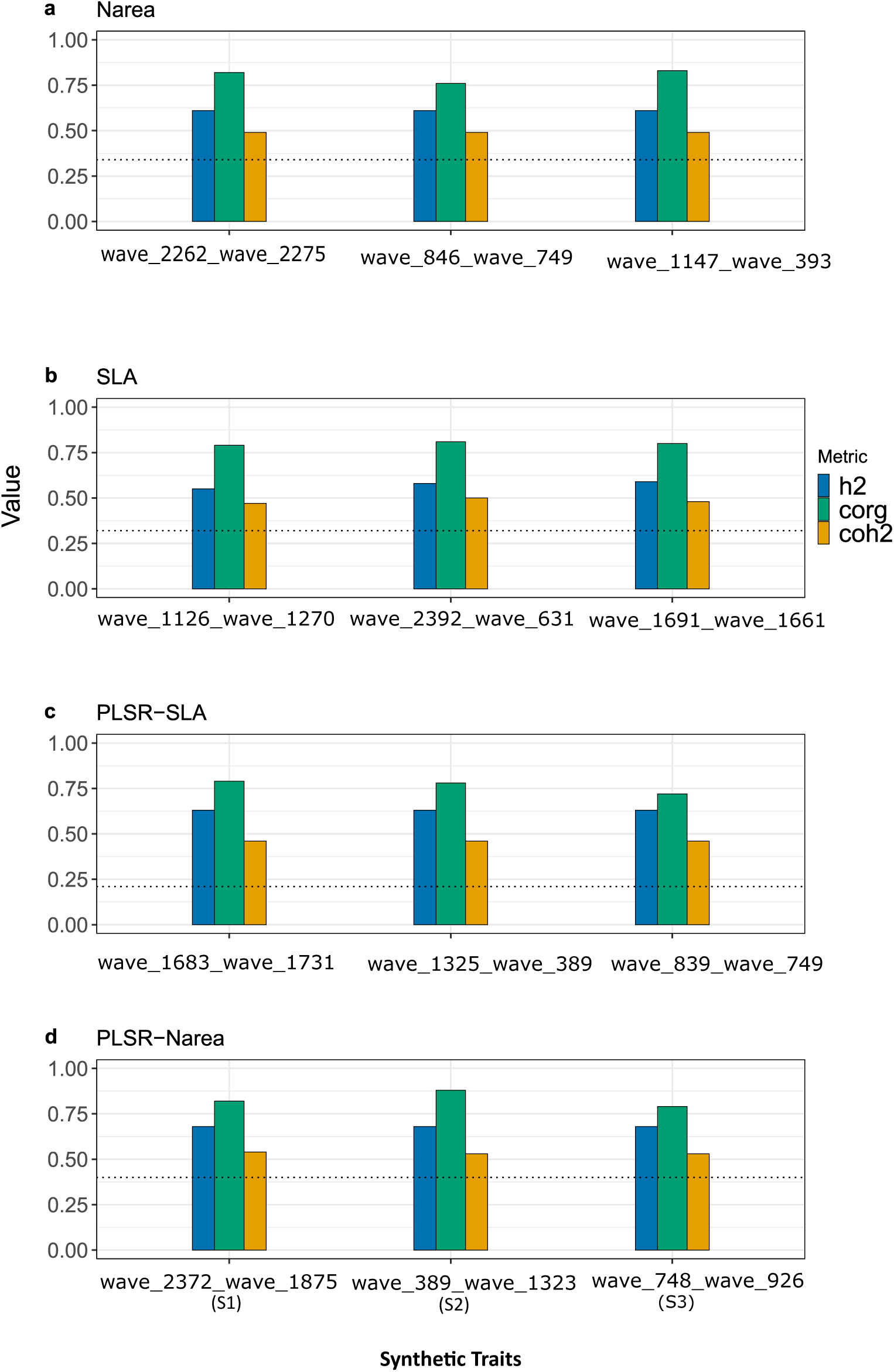
Observed heritability (*h*^2^), genetic-correlation (*cor_g_*), and co-heritability (*coh*^2^) of synthetic trait. The X-axis (Synthetic Traits) consists of S1, S2, and S3 across four target traits and twelve ratios, where three were selected for each trait, and the Y-axis consists of the value of the three different metrics observed. The target trait’s heritability is the black dotted line in each sub-figure a, b, c, and d

### 3.1 Utilizing synthetic traits in MT-GP can boost prediction accuracy compared to ST models

We selected three synthetic traits with the optimum co-*h*^2^ and used them to fit multi-trait genomic prediction models. The accuracy of the multi-trait models was consistently higher than that of the single-trait models (Figure 4). In particular, CV2 resulted in the highest prediction accuracy in all cases, whereas CV1 performed similarly to the single-trait model. Overall, the accuracy increased from 0.32 to 0.34 for Narea, 0.39 to 0.43 for SLA, 0.40 to 0.43 for PLSR-SLA, and 0.46 to 0.51 when comparing single-trait to multi-trait models. That represents an increase of approximately 6.25% for Narea (Figure 4a), 10.26% for SLA (Figure 4b), 10.87% for (Figure 4c) PLSR-Narea, and 7.5% for PLSR-SLA (Figure 4d), compared to ST.

**Figure 4.**
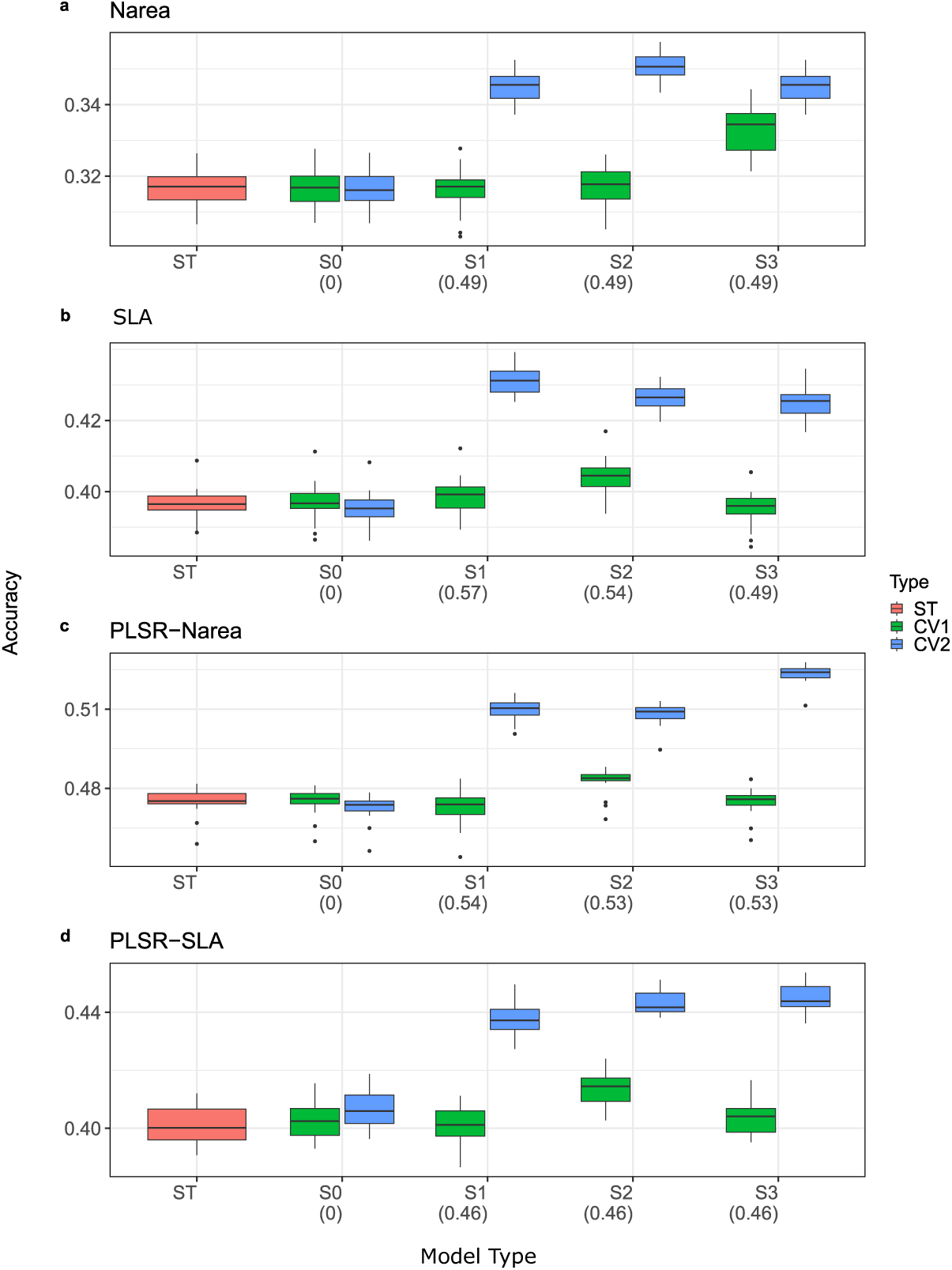
Single-trait and multi-trait genomic prediction models for four target traits: Narea, SLA, PLSR-Narea, PLSR-SLA. The Y-axis represents the prediction accuracy for multi-trait prediction utilizing synthetic traits S1, S2, and S3 in the complete dataset. The annotations on the x-axis below the synthetic trait are the co*h*^2^ value for each synthetic trait. The model was trained using EF and validated on MW. The blue boxplot is the distribution of accuracies for 20 replications for CV2, the green box plot is for CV1, and the red color is for the single-trait model.

### 3.2 Synthetic traits with lowest co-heritability did not improve prediction compared to single trait model

The idea of selecting synthetic traits based on their co*h*^2^ value was further investigated, so we fitted the model using a synthetic trait (S-0) with the lowest co*h*^2^ value to check its usefulness. Overall, the synthetic trait S-0 performed poorly in the complete model across four target traits. In general, the performance was similar to the single-trait model for both CV1 and CV2 schemes. In Figure 4 for Narea, the average accuracy of a single trait and MT was equal, i.e., 0.31, for SLA the accuracy of both ST and MT models was also equally ranged with an average accuracy of 0.39 for both models, for PLSR-SLA the accuracy of ST was 0.40 while the MT model resulted in an accuracy of 0.41, and for the accuracy of ST was 0.46. The synthetic trait with *coh*^2^ value zero selected for Narea had *h*^2^ of 0.39 and *cor_g_* of 0.02, for SLA, the *h*^2^ was 0.36, and *corg* was 0, for PLSR-Narea, the *h*^2^ was 0.43 and *cor_g_* was 0, and for PLSR-SLA the *h*^2^ was 0.51 and the *cor_g_* was 0. In summary, the factor common in all synthetic traits with *coh*^2^ zero was the lowest genetic correlation with the target trait and a low *h*^2^ value. Thus, the synthetic trait with co-heritability zero did not improve the prediction accuracy of the multi-trait model compared to the single-trait one.

### 3.3 Using only 20% of hyperspectral data is enough to improve prediction accuracy

To investigate whether or not the use of the same individuals to identify synthetic traits and to train the genomic prediction models creates a problem, we compared the accuracy trend of the complete model with five mutually exclusive subsets. The trend of prediction accuracy when utilizing only 20% of the individuals to identify the synthetic trait was very similar to the observed when utilizing the full dataset (Figure 5). In this case, the accuracy was boosted from 0.30 to 0.33 for Narea, 0.37 to 0.42 for SLA, 0.39 to 0.42 for PLSR-SLA, and 0.46 to 0.49 for PLSR-Narea across the five subsets within CV1 and CV2. In summary, the subset’s MT model prediction accuracy was increased by 18.18% for Narea, 12.2% for SLA,6.98% for PLSR-N, 4.08% for PLSR-SLA in Figure 5a, 5b, 5c and 5d compared to ST model. As expected, the numeric values of prediction accuracy were greater when utilizing the complete set of individuals, but the comparisons across models were similar.

**Figure 5.**
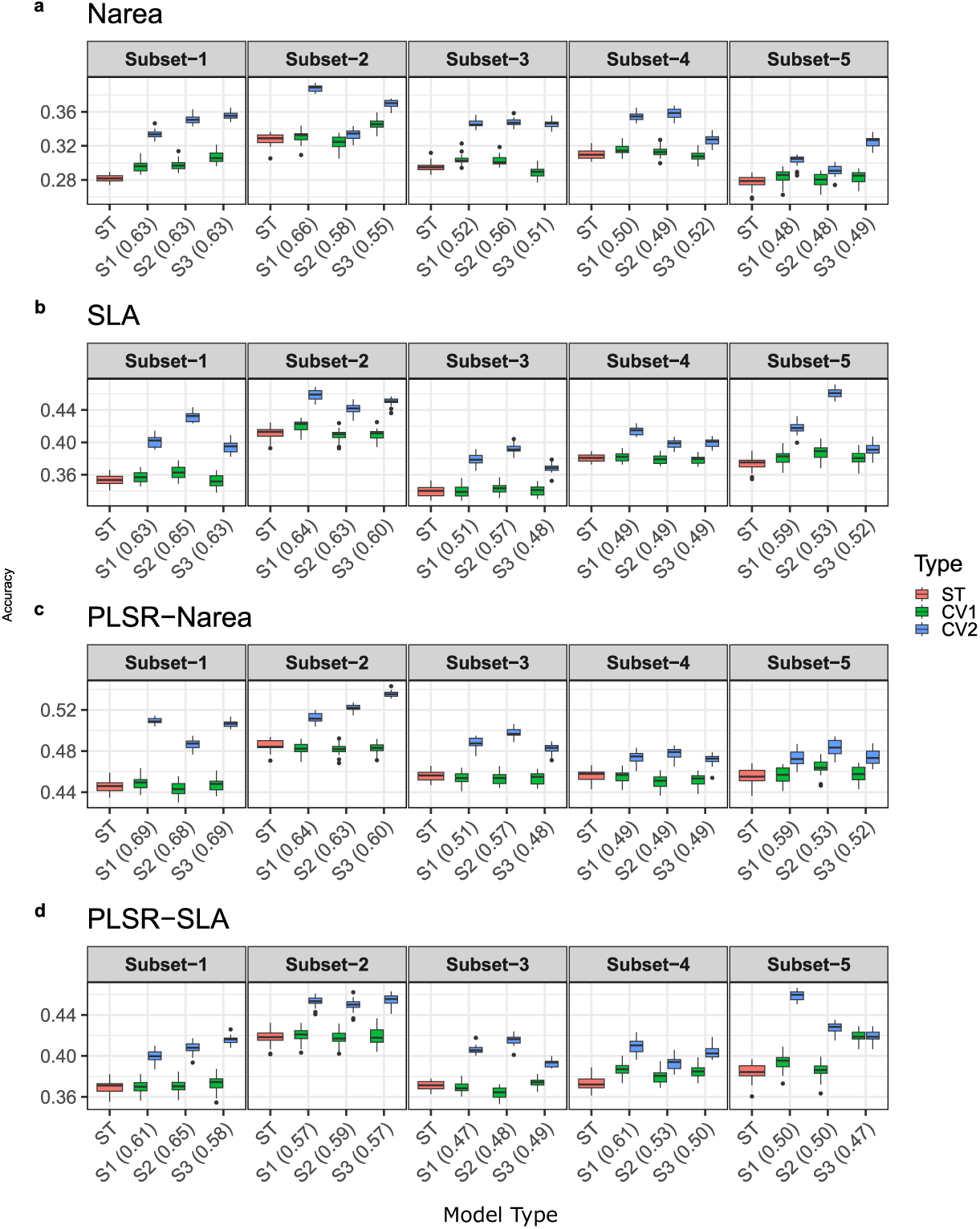
Comparison of multi-trait (MT) and single-trait (ST) models using synthetic traits identified from a subset of 20% of the individuals. The Y-axis is the accuracy of models, and the X-axis is the type of model used, i.e., MT and ST. The blue color boxplot is the distribution of accuracy of 20 replications for CV2, the green color boxplot is for CV1, and the red color is for a single trait. This data is from a model trained using EF and validated on MW.

## 4 Discussion

In this study, we generated a set of four million wavelength ratios that, as a trait set, describe a substantial fraction of the potential information that is present in a hyperspectral HTP dataset. The set of wavelength ratios was filtered using the *coh*^2^ measure to identify secondary traits for the MT-GP model. Thousands of ratios were selected within the top 1%, but since MT-GP models are computationally demanding, we selected an arbitrarily manageable number (i.e., three) of what we called synthetic traits for each target trait considered. We hypothesized that the *coh*^2^ would allow us to select synthetic traits to increase multi-trait prediction accuracy. The results supported this hypothesis, with MT-GP models consistently outperforming single-trait GP models for all four target traits. Furthermore, we demonstrated that the improvement in accuracy was only present when the synthetic trait was highly co-heritable. Overall, our analysis highlights an effective approach to leverage high-throughput phenotyping data to improve the performance of genomic prediction models.

### 4.1 High co-heritability was an indication of high prediction accuracy

Multi-trait models can leverage the correlation between traits to improve model fitting and predictive ability [41]. Furthermore, heritability has long been discussed as one of the main factors in delimiting prediction accuracy in genomic prediction [13]. In this study, we demonstrated that co-heritability can be efficiently used to select secondary traits for multi-trait prediction. The improvements observed in accuracy when utilizing synthetic traits were similar to those observed in studies utilizing traits with an intrinsic biological value as secondary traits [19, 21, 28, 36]. This illustrates the potential information hidden in HTP data. While we only picked the top three synthetic traits, this number was arbitrary, and because all of them performed similarly, we can anticipate that many other synthetic traits could be identified from the millions of wavelength ratios obtained.

Additionally, in the study, the synthetic trait with lower co-heritability, i.e., zero, which signifies either low heritability or low genetic correlation between two traits, did not show any evidence of boosting target trait predictive ability when used in the MT-GP model, which aligns with the result in the study done by [35], where using biologically meaningful secondary trait with low heritability and genetic correlation, resulted in poor performance of MT-GP model. However, despite similar *coh*^2^ values, different synthetic traits (e.g., in Narea, Figure 4 a) S1, S2, S3 have *coh*^2^ of 0.49, but they resulted in varying accuracy ranges across different MT-GP models, suggesting that further investigation on what characterizes the best secondary trait for multi-trait models is warranted.

### 4.2 Important information can be mined from hyperspectral data

Hyperspectral data has been successfully utilized in GP before by calculating vegetation indices from it [14, 40, 48]. However, we demonstrate here that only using predefined vegetation indices calculated from hyperspectral data likely fails to deliver the full potential value of hyperspectral data. Our results illustrate that a combination of wavelength ratios of high co-heritability can efficiently improve prediction accuracy regardless of being associated with a biological trait. As many indices are linear combinations of wavelength bands [4, 46], for simplicity, we opted to investigate ratios of individual wavelengths. Nonetheless, more complex spectral combinations should also be explored in the future as this method is refined.

PLSR models have proven to be useful in handling high-dimensional hyperspectral data, taking control of multi-collinearity, and providing robust, reliable measures for the prediction of traits [38, 57]. In this study, we utilized a PLSR model previously validated by Paul (2021) to predict Narea and SLA based on hyperspectral information, i.e., PLSR-Narea and PLSR-SLA. Interestingly, the heritability of the PLSR traits was slightly higher than that of the biological traits, and when combined with the synthetic trait in the multi-trait framework, the prediction accuracy for the two traits was even higher than the respective biological traits, Narea and SLA. While it is evident that any prediction model relies on ground truth data for validation, this indicates a potential for high prediction of traits such as Narea and SLA with limited requirements for slow traditional measurement methods. In addition, use of the hyperspectral data is very scalable because it allows analysis to be performed on multiple target traits (e.g., leaf N, SLA, photosynthetic capacity, sucrose content, disease status, etc) without any additional measurement effort. For plant breeding, these results reinforce the importance of HTP platforms, especially for physiologically important leaf traits that can be predicted based on spectral information.

### 4.3 Training set composition affects the MT-GP performance

The composition of the training set plays a critical role when determining the end result of many GP models [12]. The GP model is trained using the training population marker and phenotype data, which later predict the phenotype of the test set [39]. In Figure 5 (a-d), the subset-2 model consistently performed better than other subset models, regardless of the target trait we used. The sorghum diversity panel consists of different sorghum populations, which suggests the potential reason behind the subset of genotypes performing differently when used for model fitting compared to other subsets. Unlike using whole data to identify the synthetic trait and fit the GP model in a complete scenario, the subset 2 model synthetic trait was identified from random 20% of lines, and the model was fitted from the remaining 80%, where the performance of the MT-GP model using the subset-2 was better in all four traits. This suggests the effect of quality training set on the outcomes of the GP model, which can be helpful while optimizing the model [42].

Furthermore, the CV2 scheme consistently yielded better prediction accuracy than the CV1 and ST models. The CV2 model is trained by 80% of the target trait data, and 100% of hyperspectral data, where adding hyperspectral information significantly boosted the accuracy. This result is in accordance with several other instances that show that the CV2 is able to borrow information from the correlation between the secondary trait for which the additional 20% information is available [9, 19, 25].

### 4.4 Future Implications

In the past, high-throughput phenotyping data was utilized differently, such as generating vegetation indices from a few wave bands [14, 15, 30, 50], partial least square regression models [55, 59], NIRs spectroscopy and chemometrics [1] constraining the exploitation of complete information from the spectrum. However, the measurement context and the resolution of spectral data collected at different levels, i.e., leaf, plot, and field levels, can possess different applications and the nature of information [5]. We used proximally collected sensor data on the leaf level, but there is still a gap in the literature for remotely collected sensor data used in generating the secondary traits for the MT-GP model. The approach of identifying secondary traits using co-heritability measures demands further study on using field-level HTP data integrated into multiple environment multi-trait models [22], directing the future research potential in using complete spectra instead of creating ratios of wave-bands. Also, with the increase in dimension, the complexity of data increases, which suggests the potential use of different statistical and machine learning (ML) genomic prediction models that can minimize the computational time and resources to integrate HTP traits in the GP model, resulting in more confident and robust results.

## 5 Conclusions

This study demonstrated the usefulness of highly heritable correlated traits without any a priori biological value in a GP model to enhance the prediction accuracy of a target trait. The co-heritability provided us with an efficient option for selecting secondary traits, allowing the identification of helpful information within the spectra. Furthermore, the MT-GP model, powered by HTP traits, performed better than the single-trait GP model, opening up possible applications of hyperspectral-based synthetic traits for predicting complex traits.

## Acknowledgments

This research was supported by the the Artificial Intelligence for Future Agricultural Resilience, Management and Sustainability Institute (Agriculture and Food Research Initiative (AFRI) grant no. 2020-67021-32799/project accession no. 1024178 from the USDA National Institute of Food and Agriculture), Arkansas High Performance Computing Center which is funded through multiple National Science Foundation grants and the Arkansas Economic Development Commission, the Advanced Research Projects Agency-Energy (ARPA-E), U.S. Department of Energy (DOE), under Award Number DE-AR0000661, and the Office of Biological and Environmental Research in the DOE Office of Science (DE-SC0018277 and DESC0023160).

## Appendix: Supplementary File 1

**Fig1:**
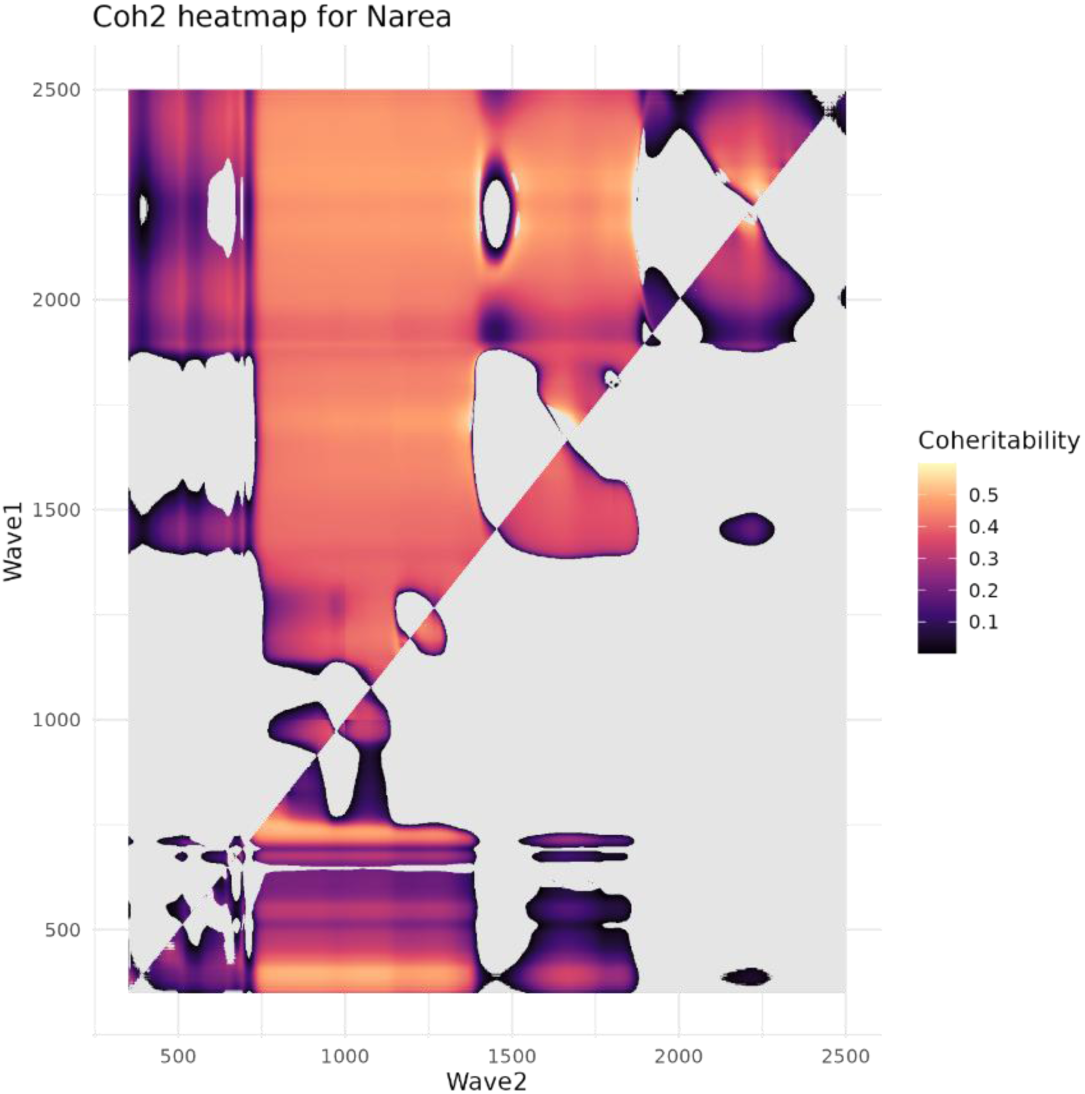
The heatmap illustrates the coheritability across and between wave-ratios for Narea, with wave1 (y-axis) and wave2 (x-axis) ranging from 350 to 2500 wavenumbers. The color gradient represents coheritability values, where darker shades indicate higher coheritability. The grey regions indicate areas where the model failed to converge or filtered during quality control.

**Fig2:**
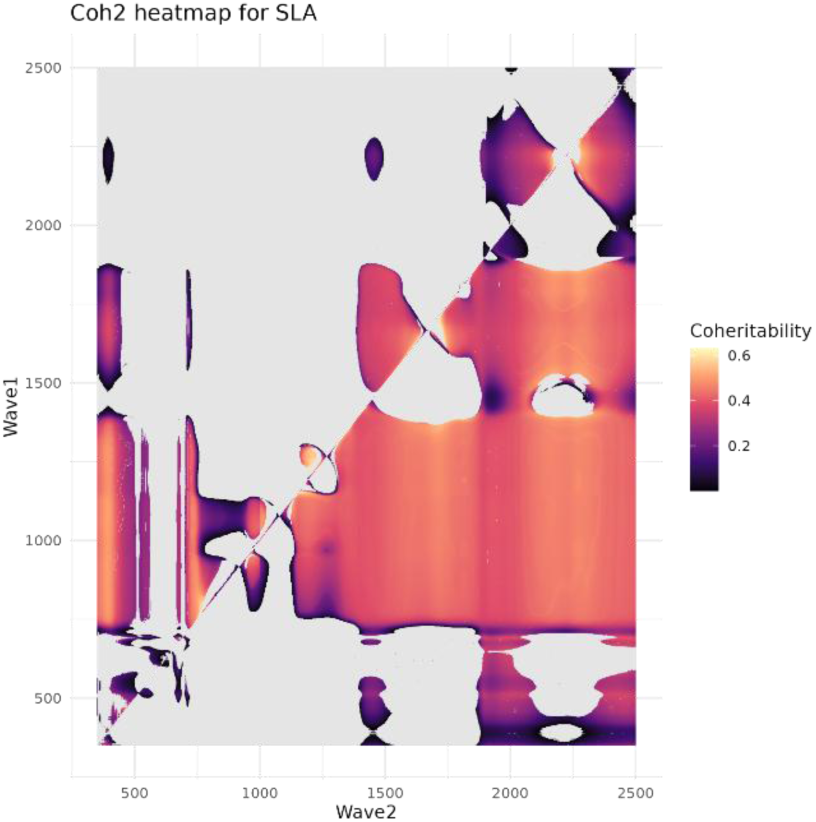
The heatmap illustrates the coheritability across and between wave-ratios for SLA, with wave1 (y-axis) and wave2 (x-axis) ranging from 350 to 2500 wavenumbers. The color gradient represents coheritability values, where darker shades indicate higher coheritability. The grey regions indicate areas where the model failed to converge or filtered during quality control.

**Fig3:**
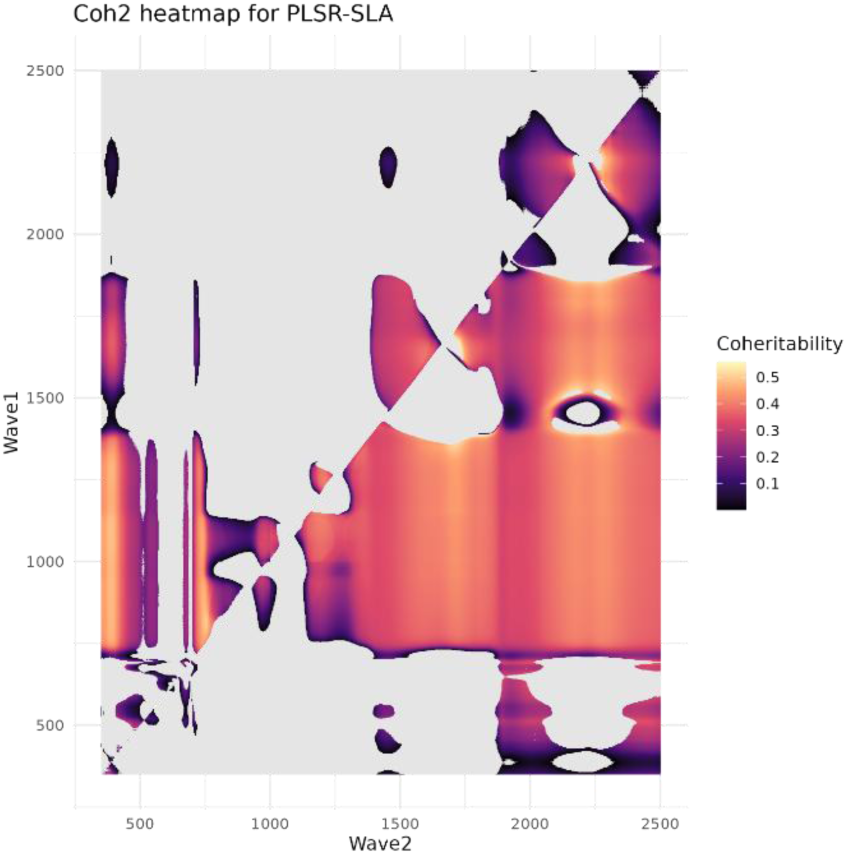
The heatmap illustrates the coheritability across and between wave-ratios for PLSR-SLA, with wave1 (y-axis) and wave2 (x-axis) ranging from 350 to 2500 wavenumbers. The color gradient represents coheritability values, where darker shades indicate higher coheritability. The grey regions indicate areas where the model failed to converge or filtered during quality control.

**Fig4:**
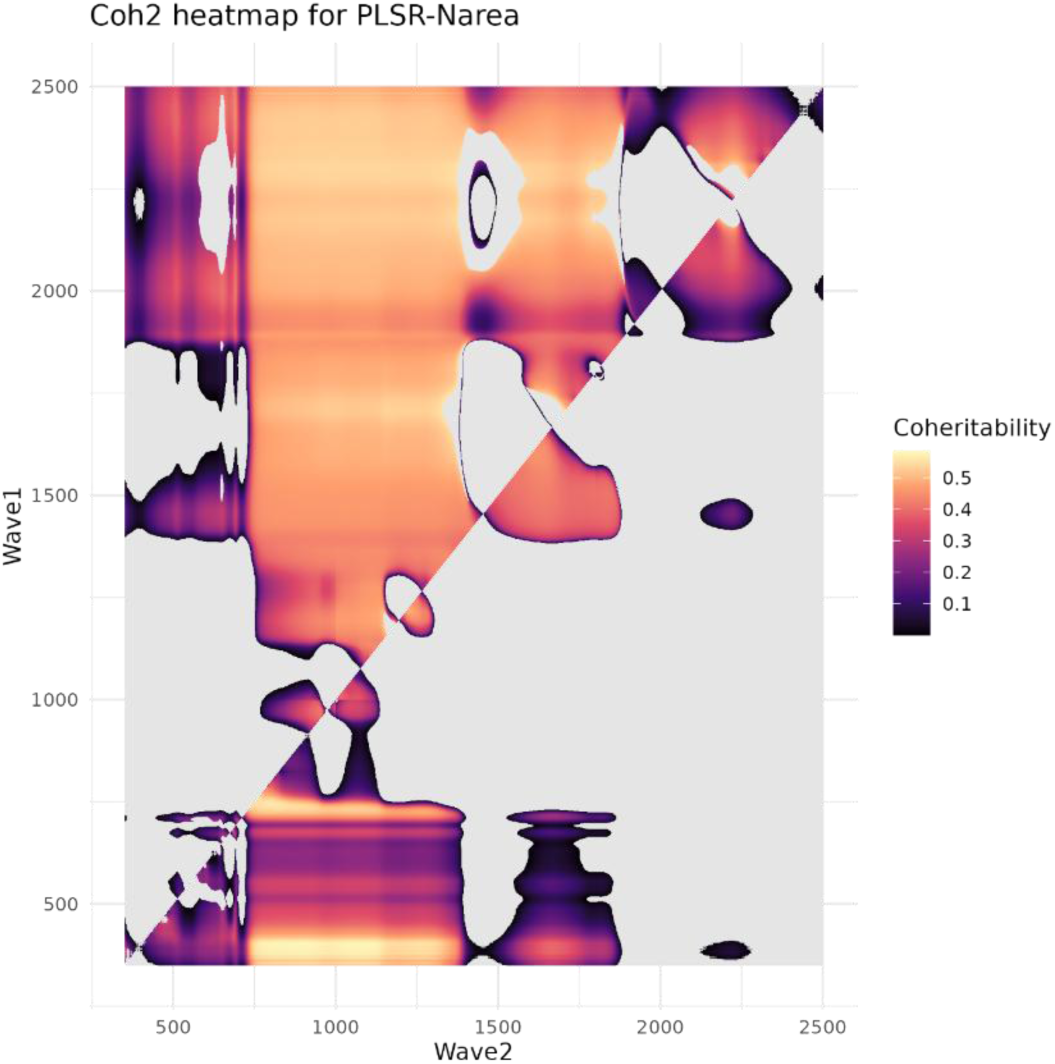
The heatmap illustrates the coheritability across and between wave-ratios for PLSR-Narea, with wave1 (y-axis) and wave2 (x-axis) ranging from 350 to 2500 wavenumbers. The color gradient represents coheritability values, where darker shades indicate higher coheritability. The grey regions indicate areas where the model failed to converge or filtered during quality control.

**Fig5:**
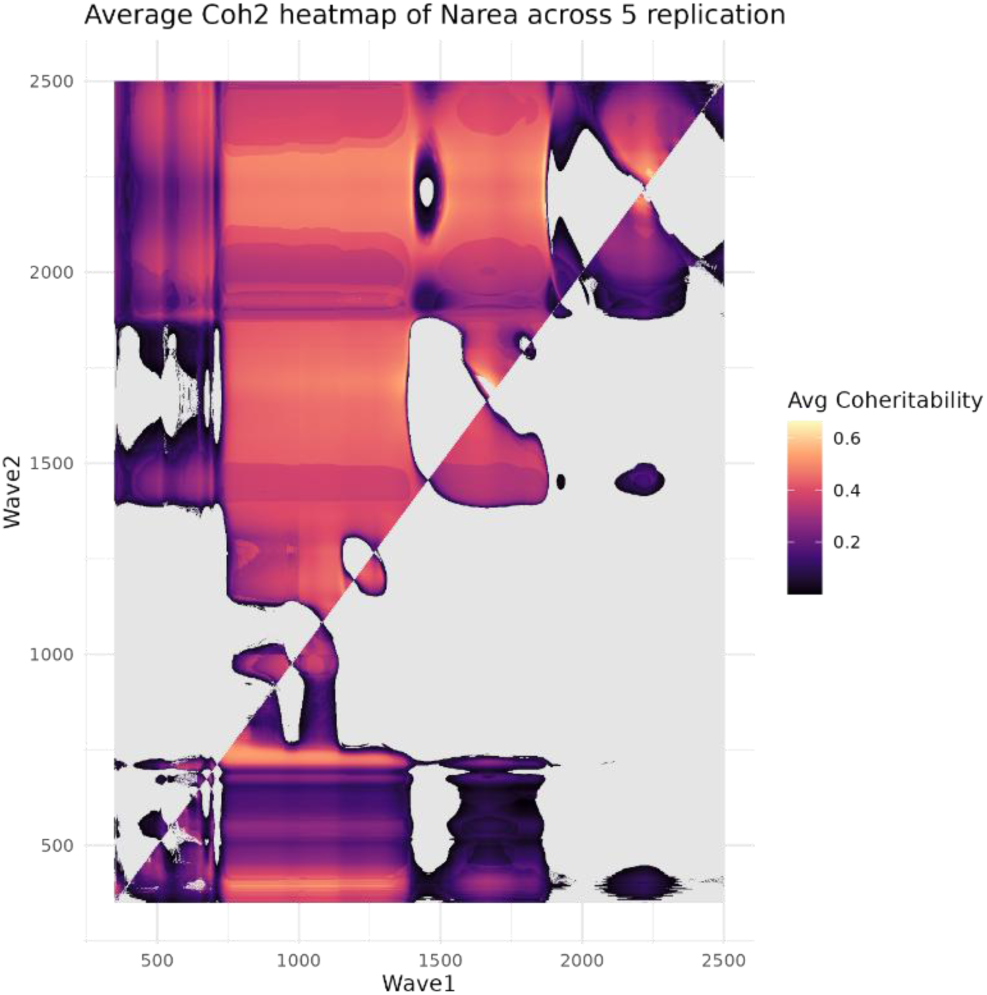
The heatmap illustrates the coheritability across and between wave-ratios for Narea averaged in all five replications, with wave1 (y-axis) and wave2 (x-axis) ranging from 350 to 2500 wavenumbers. The color gradient represents average coheritability values, where darker shades indicate higher coheritability. The grey regions indicate areas where the model failed to converge or filtered during quality control.

**Fig6:**
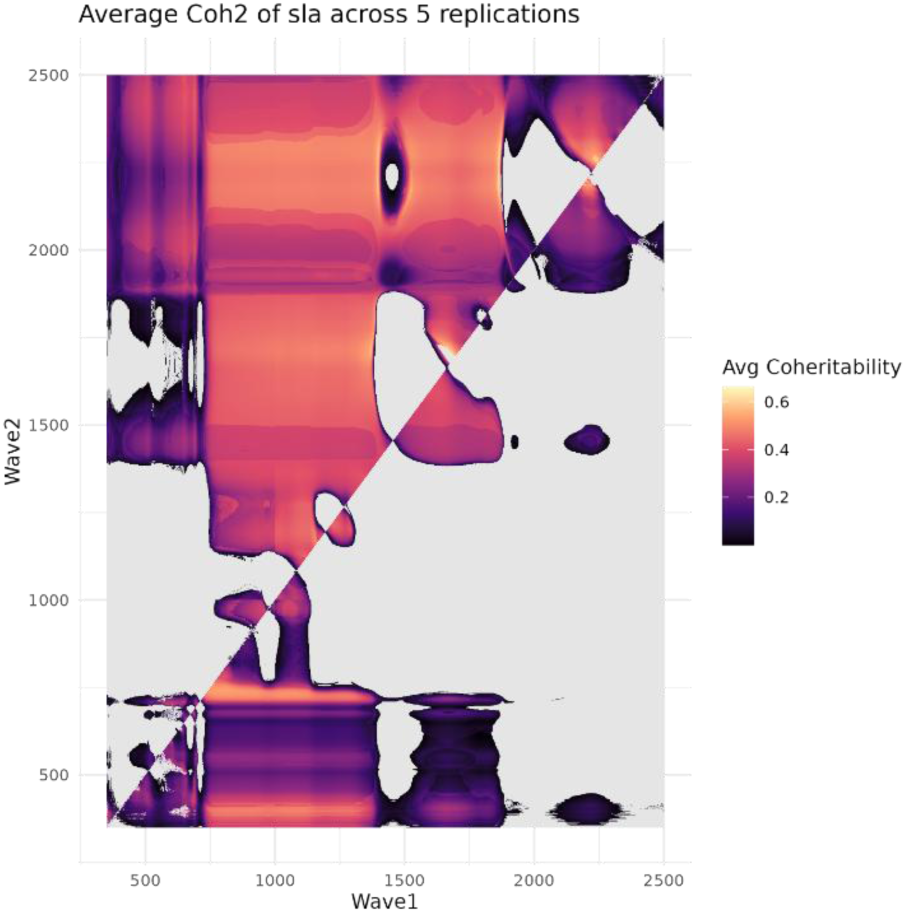
The heatmap illustrates the coheritability across and between wave-ratios for SLA averaged in all five replications, with wave1 (y-axis) and wave2 (x-axis) ranging from 350 to 2500 wavenumbers. The color gradient represents average coheritability values, where darker shades indicate higher coheritability. The grey regions indicate areas where the model failed to converge or filtered during quality control.

**Fig7:**
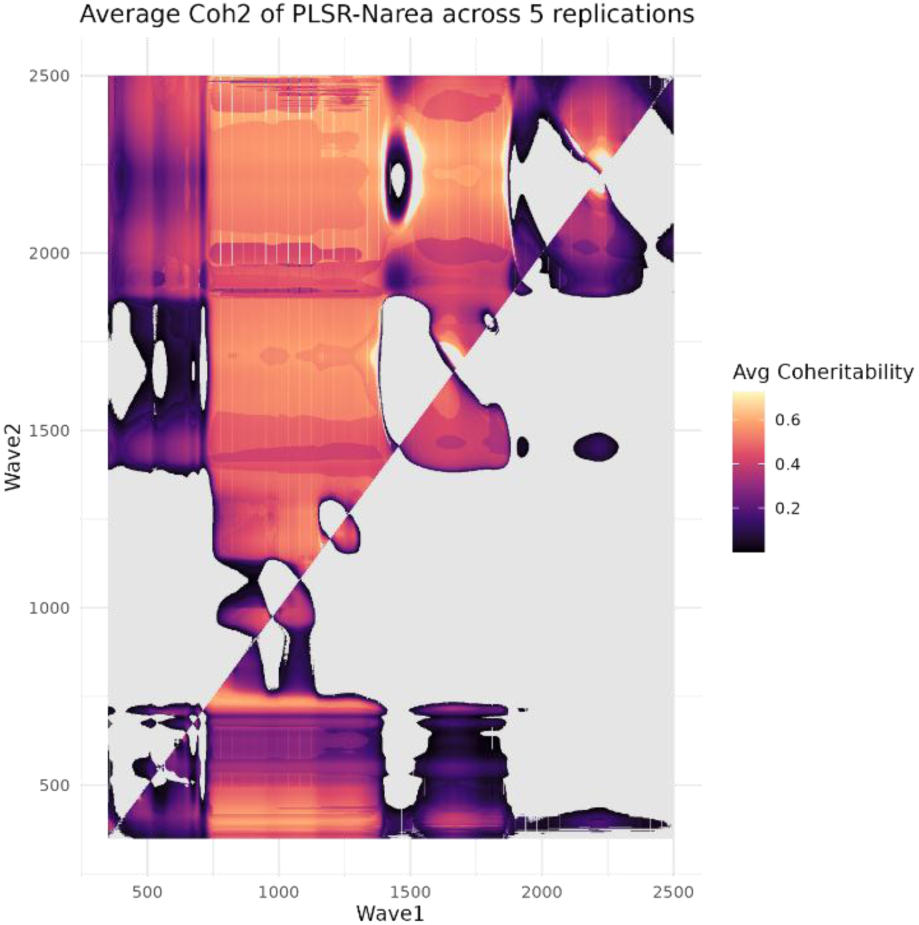
The heatmap illustrates the coheritability across and between wave-ratios for PLSR-Narea averaged in all five replications, with wave1 (y-axis) and wave2 (x-axis) ranging from 350 to 2500 wavenumbers. The color gradient represents average coheritability values, where darker shades indicate higher coheritability. The grey regions indicate areas where the model failed to converge or filtered during quality control.

**Fig8:**
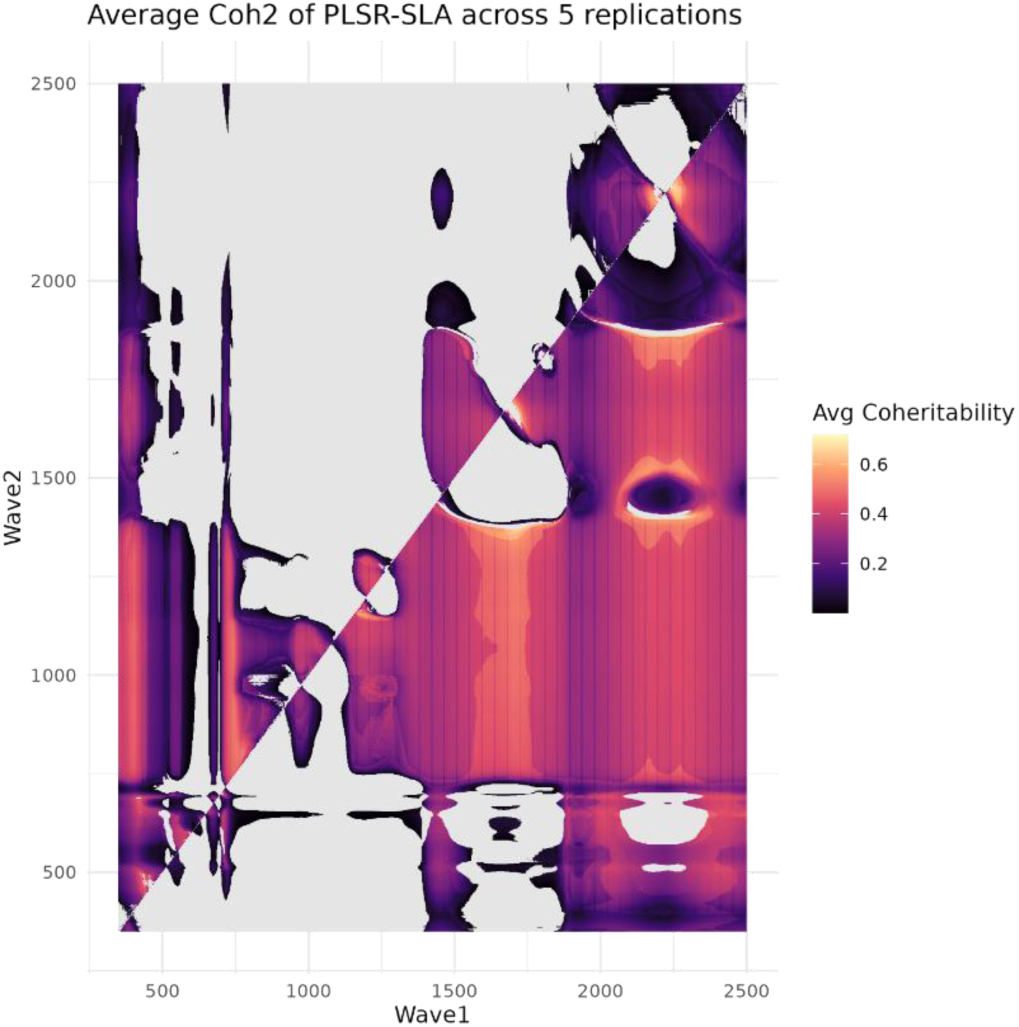
The heatmap illustrates the coheritability across and between wave-ratios for PLSR-SLA averaged in all five replications, with wave1 (y-axis) and wave2 (x-axis) ranging from 350 to 2500 wavenumbers. The color gradient represents average coheritability values, where darker shades indicate higher coheritability. The grey regions indicate areas where the model failed to converge or filtered during quality control.

